# The role of rare copy number variants in depression

**DOI:** 10.1101/378307

**Authors:** Kimberley M Kendall, Elliott Rees, Matthew Bracher-Smith, Lucy Riglin, Stanley Zammit, Michael C O’Donovan, Michael J Owen, Ian Jones, George Kirov, James T R Walters

## Abstract

The role of large, rare copy number variants (CNVs) in neurodevelopmental disorders is well-established,^1–5^ but their contribution to common psychiatric disorders, such as depression, remains unclear. We have previously shown that a substantial proportion of CNV enrichment in schizophrenia is explained by CNVs associated with neurodevelopmental disorders.^6, 7^ Depression shares genetic risk with schizophrenia^8, 9^ and is frequently comorbid with neurodevelopmental disorders^10, 11^, suggesting to us the hypothesis that if CNVs play a role in depression, neurodevelopmental CNVs are those most likely to be associated. We confirmed this in UK Biobank by showing that neurodevelopmental CNVs were associated with depression (24,575 cases, 5.87%; OR=1.36, 95% CI 1.22-1.51, p=1.61×10^-8^), whilst finding no evidence implicating other CNVs. Four individual neurodevelopmental CNVs increased risk of depression (1q21.1 duplication, PWS duplication, 16p13.11 deletion, 16p11.2 duplication). The association between neurodevelopmental CNVs and depression was partially explained by social deprivation but not by education attainment or physical illness.

---

Studies of CNVs in depression have been based on relatively small samples^12–18^, have generated inconsistent results, and no findings have met criteria for genome-wide significance. The UK Biobank offers an opportunity to investigate the relationship between CNVs and depression on a larger-scale than has hitherto been possible. We performed a CNV analysis of depression in UK Biobank data of 455,913 individuals who reported white British or Irish ethnicity (Methods). Our primary definition of depression was participant selfreport of ever having received a medical diagnosis of depression (‘self-reported depression’). To ensure our findings were not restricted to this definition, we also tested two more conservative phenotypes -(i) self-reported depression with an additional requirement for antidepressant prescription, and (ii) a hospital discharge diagnosis of depression. We sought association with depression for (i) a group of 53 CNVs known to be associated with neurodevelopmental disorders (‘neurodevelopmental CNVs’)^6, 7^ and (ii) after excluding CNVs relevant to the primary hypothesis, we tested for a residual unexplained burden among CNVs ≥ 100KB, ≥ 500KB and ≥ 1MB.

24,575 individuals (5.87%) had self-reported depression and 394,205 individuals reported no lifetime depression (Figure 1). The group of 53 neurodevelopmental CNVs was associated with depression in our primary analysis (OR=1.36, 95% CI 1.22 – 1.51, uncorrected p=1.61 x 10^-8^). Of individuals with depression 1.54% (n = 379) carried at least one of the 53 neurodevelopmental CNVs compared with 1.14% (n = 4,510) of controls. Analysis of the alternative depression phenotypes produced consistent results (Table 1), with the effect size increasing with more conservative definitions of depression. The association between the neurodevelopmental CNVs and selfreported depression remained after removing cases with a hospital discharge diagnosis of depression (OR=1.37, 95% CI 1.23 – 1.52, uncorrected p=6.27 x 10^-9^). Restricting these analyses to the subset of schizophrenia-associated CNVs generated similar results.

**Figure 1.**
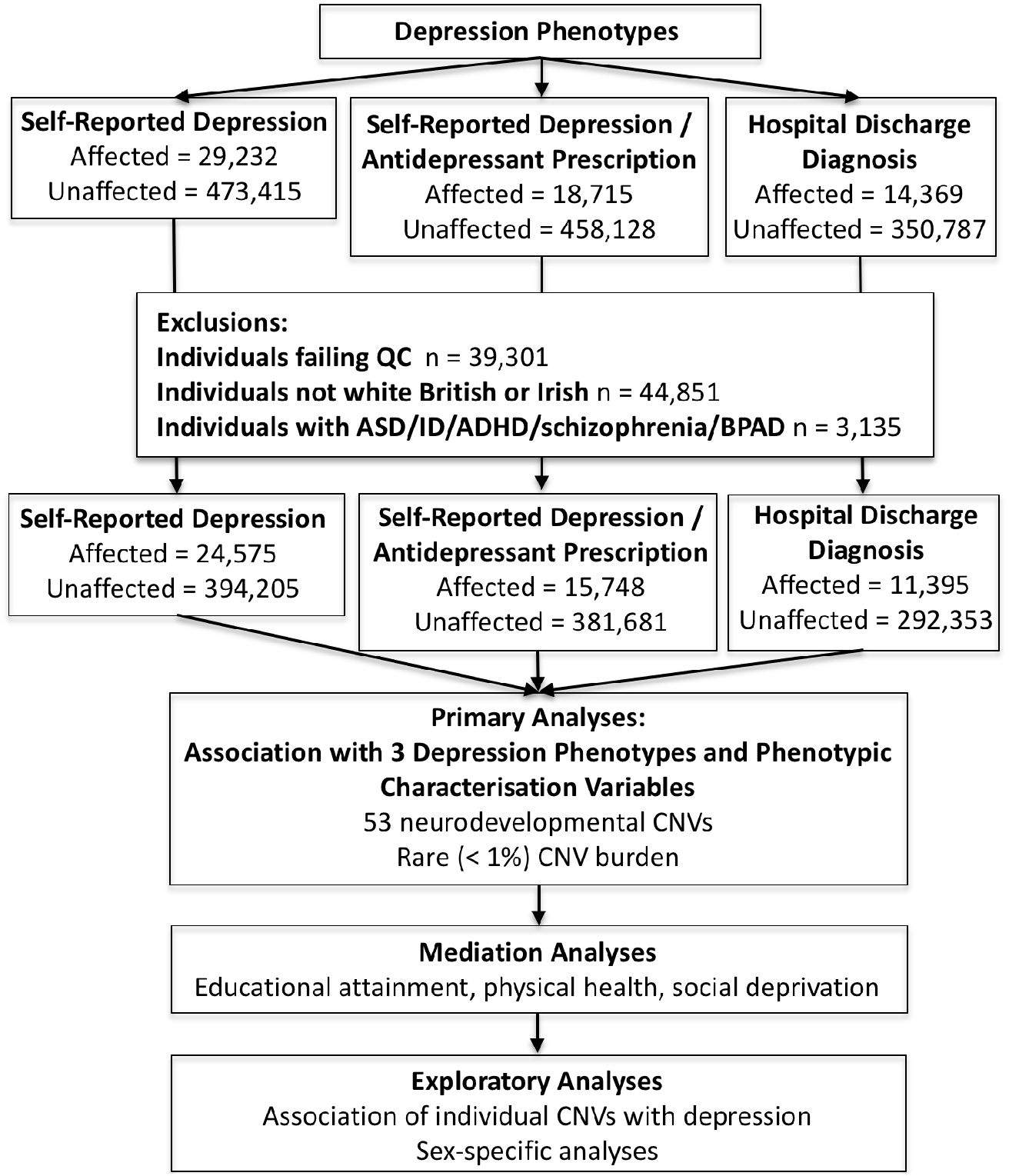
A flowchart describing the study methods.

**Table 1.** Association analyses for (i) neurodevelopmental CNVs, (ii) CNV burden with 3 depression phenotypes. Analyses were restricted to those of white British and Irish ancestry and excluded individuals with CNV-associated neurodevelopmental/neuropsychiatric disorders. The group of 53 neurodevelopmental CNVs were excluded from CNV burden analyses. OR – odds ratio, 95% CI – 95% confidence interval, p-uncorrected p value.

| CNV Type |  | Self Reported Depression<br>24,575 affected<br>394,205 unaffected |  | Self Reported Depression and<br>Antidepressant Prescription on<br>Visit 1<br>15,748 affected,<br>381,681 unaffected |  | Hospital Discharge<br>Diagnosis<br>11,395 affected<br>292,353 unaffected |  |
| --- | --- | --- | --- | --- | --- | --- | --- |
|  | Carriers | OR (95% CI) | p | OR (95% CI) | p | OR (95% CI) | p |
| Neurodevelopmental<br>CNVs | 1.2%<br>(n = 4,889) | 1.36<br>(1.22 – 1.51) | $1.61 \times 10^{-8}$ | 1.43<br>(1.26 – 1.62) | $5.31 \times 10^{-8}$ | 1.50<br>(1.29 – 1.73) | $4.07 \times 10^{-8}$ |
| Carrier Status of Rare (<1% CNVs) |  |  |  |  |  |  |  |
| 100KB + | 48.7%<br>(n = 203,853) | 1.01<br>(0.98 – 1.03) | 0.73 | 1.01<br>(0.98 – 1.04) | 0.57 | 1.04<br>(1.00 – 1.08) | 0.06 |
| 500KB+ | 8.9%<br>(n = 37,309) | 1.05<br>(1.0007 – 1.10) | 0.024 | 1.07<br>(1.02 – 1.13) | 0.01 | 1.08<br>(1.01 – 1.15) | 0.02 |
| 1MB+ | 3.4%<br>(n = 14,367) | 1.02<br>(0.95 – 1.09) | 0.59 | 1.03<br>(0.95 – 1.13) | 0.45 | 1.03<br>(0.94 – 1.14) | 0.52 |

After exclusion of the 53 neurodevelopmental CNVs, there was a weak association between CNVs ≥ 500KB and depression, which did not survive correction for multiple testing. There was no evidence for association between CNVs ≥ 100KB, ≥ 1MB and depression (Table 1).

Exploratory analyses of individual neurodevelopmental CNVs found 10 CNVs to be nominally associated with self-reported depression, of which four (1q21.1 duplication, Prader Willi syndrome duplication, 16p13.11 deletion and 16p11.2 duplication) survived Bonferroni correction for the 53 CNVs tested (p value threshold 0.00094, Figure 2, Supplementary Table 1).

**Figure 2.**
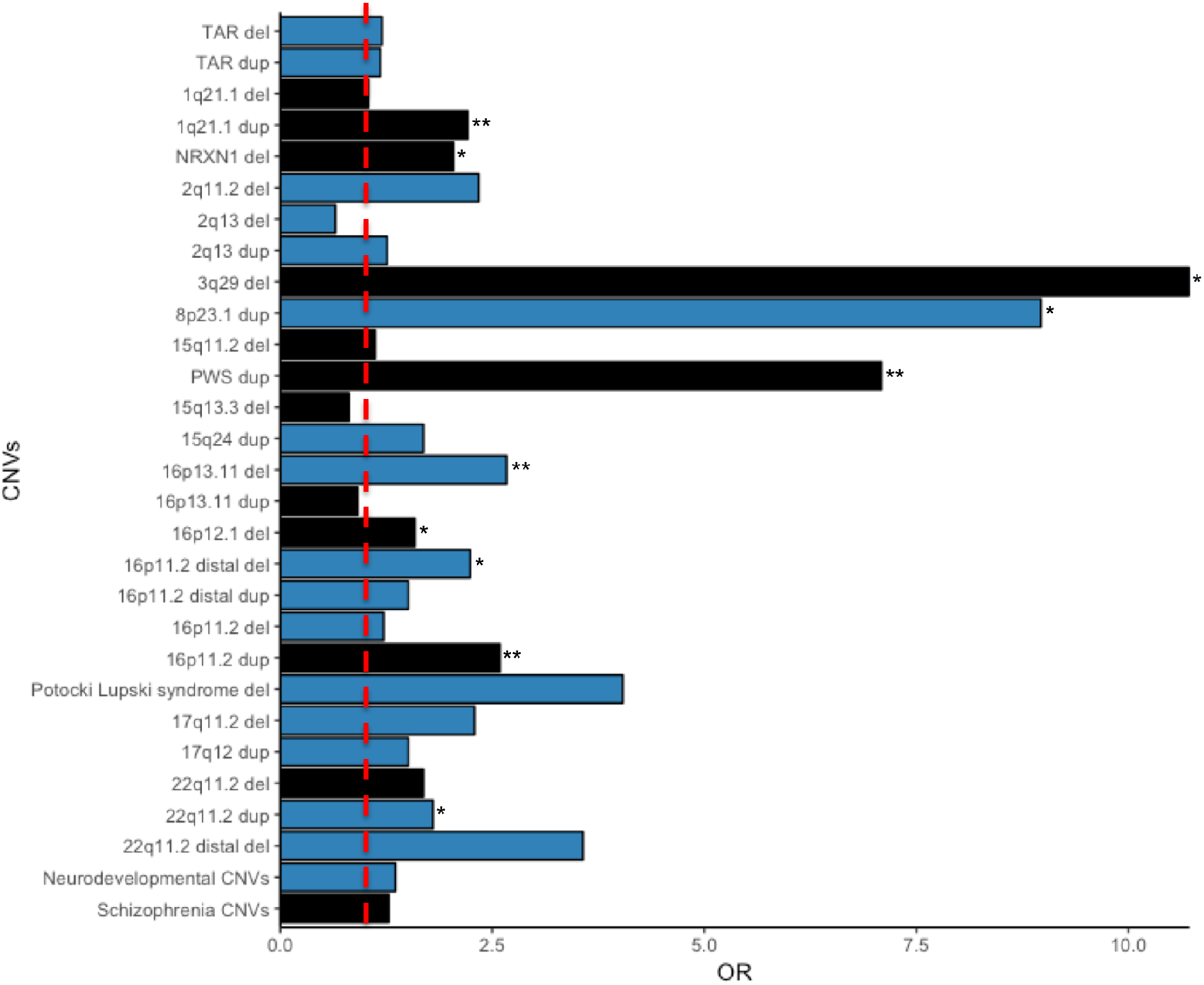
Analyses of individual neurodevelopmental CNVs for association with self-reported depression. Results with a p value<0.05 are marked with *, with those surviving Bonferroni correction for 53 tests being marked with **. Schizophrenia CNVs are coloured black. All the CNVs fall within the neurodevelopmental CNV group. The broken red line indicates an OR (odds ratio) of 1.

The addition of genotyping platform as a covariate in regression analyses did not change the results as presented.

We used data from 157,397 individuals who completed an online follow-up mental health questionnaire to examine association between CNVs and markers of depression severity (age at onset, number of episodes of depression, duration of worst depressive episode). We restricted these analyses to the 73,234 individuals who reported experience of prolonged feelings of sadness or depression (61,467 unaffected), a phenotype which itself was associated with neurodevelopmental CNV carrier status (OR=1.19, 95% 1.06-1.33, p=0.004). We did not find any association between neurodevelopmental CNVs and markers of depression severity that survived correction for multiple testing (Supplementary Table 2).

In order to better understand the relationship between neurodevelopmental CNVs and depression, we investigated whether the association was explained by measures of (i) educational attainment (qualifications), (ii) physical health (number of non-psychiatric hospital admission diagnoses) and (iii) social deprivation (Townsend Deprivation Index). These variables were chosen because of their known associations with depression^19, 20^, their postulated or proven association with CNVs and their availability in a large proportion of the UK Biobank (UKBB) sample. The association between neurodevelopmental CNVs and depression was partially explained by social deprivation (Figure 3, Supplementary Tables 3a, 3b) but not by level of education or physical health. There remained a strong independent association between these CNVs and depression in analyses incorporating these other measures.

**Figure 3.**
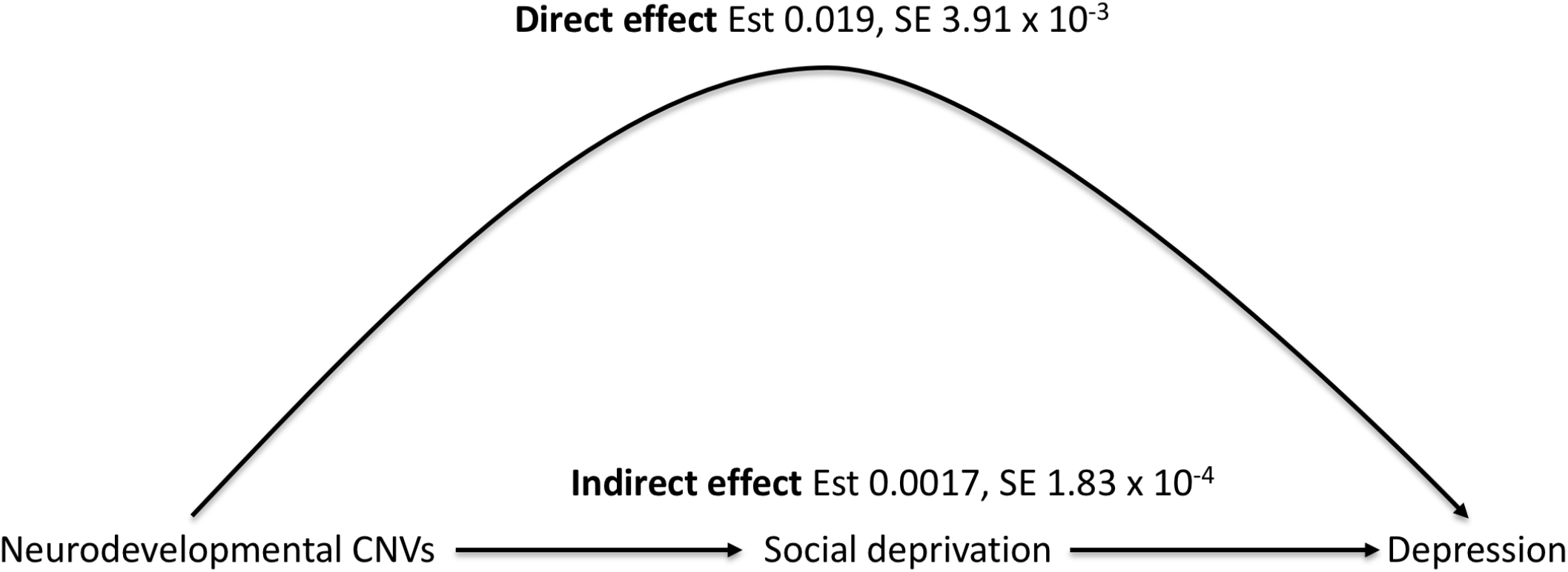
Further investigation of the association between neurodevelopmental CNVs and depression via social deprivation (educational attainment and physical health results are shown in Supplementary Tables 3a and 3b]. Est: Estimate of effect size, SE: standard error of estimate

Given recent evidence for an increased rate of large CNVs in female children with anxiety or depression^21^, we undertook exploratory analyses which provided weak evidence for a higher rate of depression in female neurodevelopmental CNV carriers than males. This increased rate was over and above the baseline increased rate of self-reported depression in females (interaction term OR=0.78, 95% CI 0.62 – 0.98, uncorrected p=0.03, Supplementary Table 4) although this effect was weaker still and non-significant for our secondary depression definitions.

Here we report the largest study of CNVs in depression to date. We tested two main hypotheses - (i) that risk CNVs for neurodevelopmental disorders would be associated with depression and, (ii) that, after excluding CNVs relevant to the primary hypothesis, there would be a residual unexplained burden among CNVs ≥ 100KB, ≥ 500KB and ≥ 1MB. Our results support our first hypothesis - CNVs previously associated with neurodevelopmental disorders were associated with increased risk of lifetime depression, whether defined on the basis of selfreported diagnosis, this combined with antidepressant treatment or on hospital discharge diagnosis. Four neurodevelopmental CNVs (1q21.1 duplication, PWS duplication, 16p13.11 deletion, 16p11.2 duplication) were individually associated with depression at levels of significance that survived Bonferroni correction for the 53 CNVs tested. We note that none of these CNV loci overlap with risk loci recently identified in a large depression genome-wide association study^9^. The risk of depression in CNV carriers in this study (whether CNVs are considered individually or collectively) was lower than that identified in previous studies of schizophrenia but qualitatively, the results followed a similar pattern - the highest risk for both disorders was conferred by 3q29del (depression OR=10.72, schizophrenia OR =57.65) and the lowest risk for both disorders was conferred by 16p12.1del (depression OR=1.59, schizophrenia OR=3.3).^5, 6^ After excluding neurodevelopmental CNVs we found no evidence of residual burden of risk for depression among CNVs ≥ 100KB, ≥ 500KB and ≥ 1MB.

Further investigation of the association between neurodevelopmental CNVs and depression indicated that this relationship is partially explained by social deprivation. To our knowledge this is the first report showing these CNVs are associated with neighbourhood measures of social deprivation, and thus implicates an important mechanism by which CNV carrier status could increase risk of depression, although longitudinal data on these measures is needed to establish the causal directionality between depression and social deprivation.

We acknowledge some limitations of this study. Our primary depression definition relied on self-report, a method known to be subject to information bias.^22^ However, this is unlikely to have markedly influenced our findings given the almost identical, or even stronger, results from using the clinicians’ hospital discharge diagnosis of depression phenotype. Another limitation is the relatively low rate of depression compared to population estimates^23^. While this likely reflects the better than average health and functioning of the UK Biobank sample, and imprecision in the definition of disorder, these factors are not likely to have generated spurious CNV associations.

Neurodevelopmental CNVs have incomplete penetrance for major developmental disorders^24^, yet beyond mild cognitive impairment^25, 26^, little is known about phenotypic associations with CNV carrier status. Our study is the first to robustly demonstrate association between these CNVs and depression and thus extends the spectrum of clinical phenotypes that are associated with CNV carrier status.

## Methods

### Sample

Between 2006 and 2010, UK Biobank recruited ~500,000 individuals (54% female) aged 37 – 73 years living in the United Kingdom. Ethical approval was granted to UK Biobank by the North West Multi-Centre Ethics Committee and all participants provided informed consent to participate in UK Biobank projects. Phenotypic data were collected at assessment centres via touchscreen devices and nurse-led interviews. Participants provided blood, urine and saliva samples. This study was conducted under the conditions of UK Biobank project number 14421.

### Depression Phenotypes

Analyses of depression in the UK Biobank to date have used multiple definitions of the disorder.^27, 28^ In view of this lack of consensus regarding case definition, we began by using a relatively liberal definition of lifetime depression, rating as cases those individuals who reported a doctor had told them they have depression (self-reported depression code 1286, UKBB field 20002. We repeated our analyses using two alternative, more conservative definitions of depression – lifetime self-reported depression with current antidepressant prescription at the time of visit 1 and, hospital discharge diagnosis of depression. *Self-reported depression with antidepressant prescription at visit 1 -* we constructed a binary depression variable using (i) the self-reported depression code 1286 in UKBB field 20002 and (ii) antidepressant prescription codes in UKBB field 20003. Individuals were included as affected if they reported that a doctor had told them they have depression and they were prescribed an antidepressant medication at the time of first assessment. Individuals, who fulfilled only one of the two criteria i.e. self-reported depression or antidepressant prescription alone, were excluded from analyses. *Hospital discharge diagnosis of depression* – individuals were included as affected if they had a hospital admission with a primary or secondary ICD-10 code for depression (UKBB fields 41202 and 41204) (Figure 1). For individuals assessed at Scottish assessment centres, hospital records covered general hospitals but not psychiatric hospitals (for Wales and England these covered both). For these individuals, we accepted a secondary ICD-10 code for depression as evidence of depression diagnosis as this could be coded during admission to a general hospital. However, in the absence of primary ICD-10 codes for depression, we were unable to determine the absence of depression in controls. Therefore, we removed Scottish controls from this variable.

In 2017, 157,397 individuals completed an online follow-up mental health questionnaire. We aimed to further characterise associations with depression phenotypes in individuals who stated that they had ever experienced prolonged feelings of sadness or depression on this questionnaire (UKBB field 20446). We examined data from the following variables - (i) age at first episode of depression (UKBB field 20433), (ii) duration of worst depression (UKBB field 20438) and (iii) lifetime number of depressed episodes (UKBB field 20442). *Duration of worst depression (UKBB field 20438)* - this was coded in ranges of months e.g. less than a month, between one month and three months. Previous data has shown that the median duration of a depressive episode is 3 months.^29^ Therefore, this variable was dichotomised into 0-3 months and more than 3 months. *Lifetime number of depressed episodes (UKBB field 20442)* - this variable was dichotomised using a median split approach (median = 1).

### Genotyping and CNV Calling

DNA was extracted from whole blood^30^ and then genotyped at the Affymetrix Research Services Laboratory, Santa Clara, CA on the UK Biobank Axiom and UK BiLEVE arrays. Genotypes were released to Cardiff University after application to UK Biobank (project number 14421). CNV calling was carried out using PennCNV-Affy^31^ using biallelic markers common to both genotyping platforms and is described in detail elsewhere.^25^ CNV burden analysis was carried out on the CNV calls generated by Penn-CNV-Affy using PLINK 1.07.^32^ Individual samples were excluded if they had ≥ 30 CNVs, a waviness factor of >0.03 or < 0. 03, a SNP call rate of <96% or LRR SD of >0.35. Individual CNVs were excluded if they were covered by <20 probes, had a density coverage of <1 probe per 20,000 base pairs or a confidence score of <10.

### Defining Copy Number Variant Sets and Analysis

Following the approach of our recent study using UK Biobank data^25^ we defined a group of ‘neurodevelopmental CNVs’ as those 54 CNVs for which there is at least nominally significant evidence for association with neurodevelopmental disorders(p < 0.05).^3^ We excluded the high frequency 15q11.2 duplication resulting in a final list of 53 ‘neurodevelopmental CNVs’. In exploratory analyses each of the CNVs for which there were ≥5 observations were analysed individually for association with self-reported depression.

CNV burden analyses were carried out using PLINK on regions of variable copy number at three size thresholds: (i) ≥ 100KB, (ii) ≥ 500KB and (iii) ≥ 1MB. CNVs were filtered for frequency at <1% using the --cnv-freq-exclude-above command and overlapping lower copy repeat regions were filtered out using the --cnv-exclude command. PLINK outputs were converted into CNV carrier status, which was used as the outcome in regression analyses. For all burden analyses, the group of 53 CNVs associated with neurodevelopmental disorders were excluded.

Association analyses were carried out in R using logistic or linear regression as appropriate with age and sex as covariates. Analyses were restricted to those who reported white British or Irish ethnicity and we excluded individuals with CNV-associated neurodevelopmental/neuropsychiatric disorders (autism spectrum disorder (ASD), intellectual disability (ID), attention deficit hyperactivity disorder (ADHD), schizophrenia or bipolar affective disorder (BPAD)) from being cases and controls.

### Further Investigation of the Neurodevelopmental CNV-Depression Association

In order to better understand the association between neurodevelopmental CNVs and depression we investigated whether the association was explained by three variables known to be associated with depression^19, 20^ and postulated to be associated with CNVs – (i) educational attainment (qualifications), (ii) physical health (number of non-psychiatric hospital admission diagnoses) and, (iii) social deprivation (Townsend Deprivation Index). *Educational attainment:* Prior to analysis, data from the academic qualifications field were dichotomised and recoded into college/university degree or all other qualifications, an approach previously used for this data field (UKBB field 6138).^33^ *Physical health:* We used number of hospital admission physical health codes as a proxy measure of the extent of physical health problems (excluding psychiatric codes; UKBB fields 41202 and 41204). *Social deprivation:* This was measured using Townsend Deprivation Index codes (UKBB field 189). Analyses were carried out using structural equation modelling in the lavaan package in R.^34^ The proportion explained was estimated by indirect effect/total effect. Age and sex were used as covariates throughout the analyses.

### Sex-Specific Analyses

Previous studies have reported a small but significant excess of large (≥ 500KB) rare (< 1%) CNVs in females.^35^ Recent evidence has also suggested that female children with anxiety or depression are more likely to carry large CNVs than males.^21^ This led us to examine rates of depression in female and male CNV carriers in our sample. Following finding that there was an excess of female CNV carriers with depression, we added an interaction term consisting of the product of neurodevelopmental CNVs and sex, to our main regression model.

## Acknowledgements

This work was funded by a Wellcome Trust Clinical Research Training Fellowship awarded to K Kendall. The work at Cardiff University was supported by Medical Research Council (MRC) Centre (MR/L010305/1), Program (G0800509) and Project (MR/L011794/1) grants.

